# X-ray and solution structures of human beta-2 glycoprotein I reveal a new mechanism of autoantibody recognition

**DOI:** 10.1101/2020.02.25.963314

**Authors:** Eliza Ruben, William Planer, Mathivanan Chinnaraj, Zhiwei Chen, Xiaobing Zuo, Vittorio Pengo, Vincenzo De Filippis, Ravi K. Alluri, Keith R. McCrae, Paolo Macor, Francesco Tedesco, Nicola Pozzi

**Affiliations:** Edward A. Doisy Department of Biochemistry and Molecular Biology, Saint Louis University School of Medicine, St. Louis, MO 63104; X-Ray Science Division, Argonne National Laboratory, Argonne, Illinois 60439; Division of Cardiac, Thoracic, and Vascular Sciences, University of Padova, Padova, Italy; Department of Pharmaceutical & Pharmacological Sciences, University of Padua, via F. Marzolo 5, 35131 Padua; Department of Cellular and Molecular Medicine, Cleveland Clinic, Cleveland, OH, United States; Department of Life Sciences, University of Trieste, Italy; Istituto Auxologico Italiano, IRCCS, Laboratory of Immuno-Rheumatology, Milan, Italy.

## Abstract

Venous and arterial thromboses in patients suffering from the autoimmune disorder Antiphospholipid Syndrome (APS) are caused by the presence of antiphospholipid antibodies (aPL). Emerging evidence indicates that autoantibodies targeting the epitope R39-R43 in the N-terminal domain, Domain I (DI), of β_2_-glycoprotein I (β_2_GPI) are among the most pathogenic aPL in patients with APS. How such autoantibodies engage β_2_GPI at the molecular level remains incompletely understood. Here, we have used X-ray crystallography, single-molecule FRET, and small-angle X-ray scattering to demonstrate that, in the free form, under physiological pH and salt concentrations, human recombinant β_2_GPI adopts an elongated, flexible conformation in which DI is exposed to the solvent, thus available for autoantibody recognition. Consistent with this structural model, binding and mutagenesis studies revealed that the elongated form interacts with a pathogenic anti-DI antibody in solution, without the need of phospholipids. Furthermore, complex formation was affected neither by the neighboring domains, nor by the presence of the linkers, nor by the glycosylations. Since the pathogenic autoantibody requires residues R39 and R43 for optimal binding, these findings challenge longstanding postulates in the field envisioning β_2_GPI adopting immunologic inert conformations featuring inaccessibility of the epitope R39-R43 in DI and support an alternative model whereby the preferential binding of anti-DI antibodies towards phospholipid-bound β_2_GPI arises from the ability of the pre-existing elongated form to bind to the membranes and then oligomerize, processes that are likely to be supported by protein conformational changes. Interfering with these steps may limit the pathogenic effects of anti-DI antibodies in APS patients.

**Significance:** In the autoimmune disorder called Antiphospholipid Syndrome (APS), the presence of autoantibodies targeting the plasma glycoprotein beta-2 glycoprotein I (β_2_GPI) is associated with arterial and venous thrombosis as well as pregnancy complications. Understanding how β_2_GPI becomes immunogenic and how autoantibodies in complex with β_2_GPI cause the blood to clot remains a top priority in the field. By elucidating the structural architecture of β_2_GPI free in solution, our studies challenge longstanding postulates in the field and shed new light on the pathogenic mechanisms of APS that may help the development of new diagnostics and therapeutic approaches.

## Introduction

β_2_GPI is a 50-kDa multi-domain glycoprotein that circulates in the plasma at a concentration of 0.2 mg/ml(1, 2)(Fig 1A). It acquired centerstage in hematology in 1990 when it was recognized by two independent studies as the dominant antigen of antiphospholipid antibodies (aPL) in the Antiphospholipid Syndrome (APS)(3–5), a life-threatening blood clotting disorder characterized by vascular thrombosis and pregnancy morbidity(6). Autoantibodies against β_2_GPI (anti-β_2_GPI) are indeed frequently found in young patients with a history of thrombosis(7, 8); they are often associated with lupus anticoagulant, a laboratory test that indicates predisposition for blood clots(9); they induce(10) and potentiate thrombus formation *in vivo* (11, 12) and cause pregnancy complications resulting in fetal loss(13).

**Figure 1.**
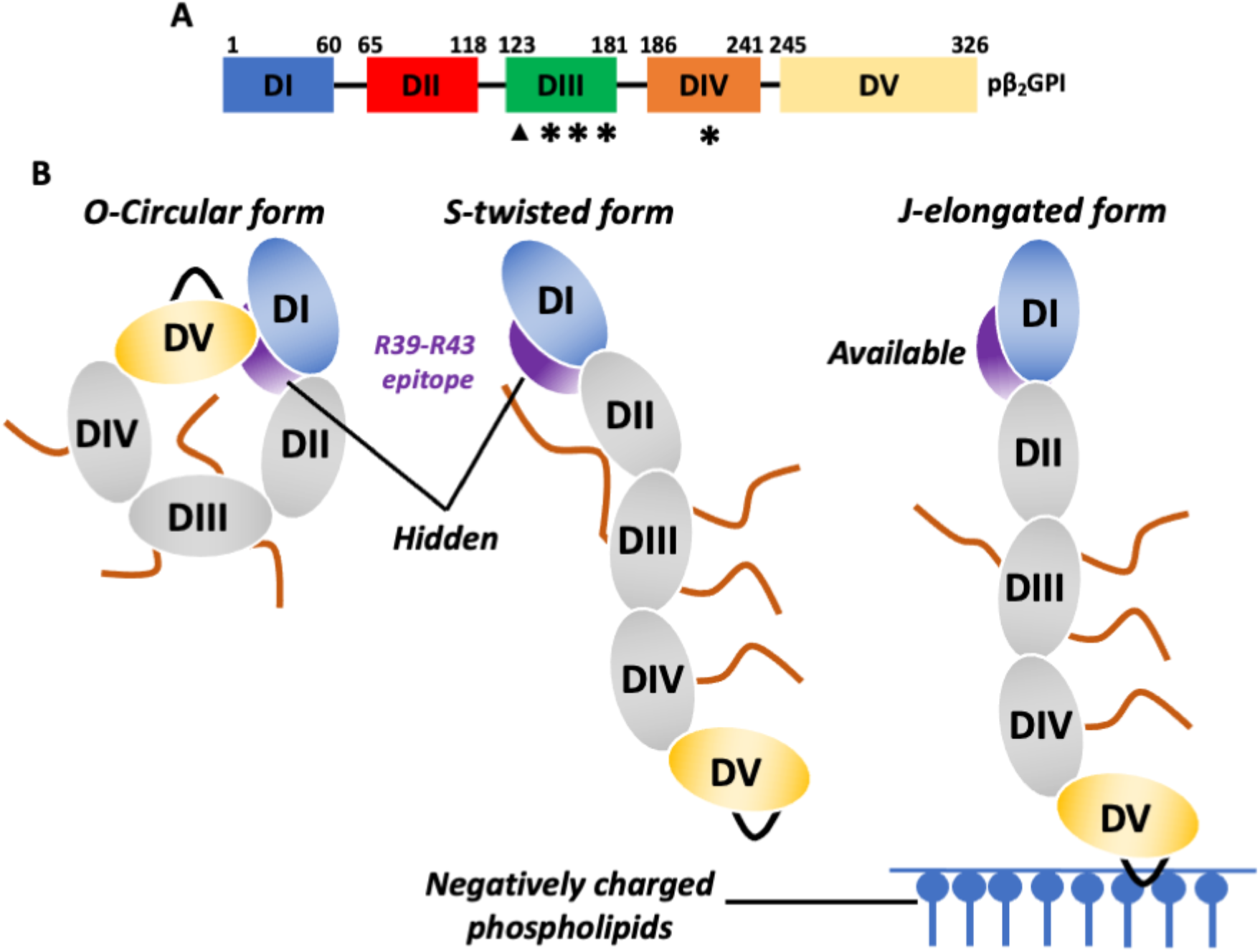
Structure of human β_2_GPI and current mechanism of antigen-antibody recognition for pathogenic anti-DI antibodies in APS. **(A)** Color-coded domain structure of human β_2_GPI (pβ_2_GPI). β_2_GPI consists of 326 amino acids organized into five domains (DI-V) connected by four short linkers, Lnk1 (residues 61-64); Lnk2 (residues 119-122), Lnk3 (residues 182-185) and Lnk4 (residues 242-244)(64). Domains I-IV are canonical complement control protein (CCP) domains, each containing two disulfide bonds. In contrast, DV is aberrant, consisting of one extra disulfide bond and a 19-residue hydrophobic loop that is responsible for anchoring the protein to negatively charged phospholipids(47, 65). Similar to other CCP-containing proteins, β_2_GPI is also heavily glycosylated, bearing four N-linked and one O-linked glycosylations located at positions T130, N143, N164, N174, and N234 that account for ~20% of the total protein mass. The position of the O- and N-linked glycosylations is shown as triangles (▲) and stars (*), respectively. **(B)** Based on previous studies, β_2_GPI is believed to adopt an O-circular(27, 28), an S-twisted(26) and a J-elongated conformation(24, 25). The J-open conformation results upon interaction of DV with the phospholipids exposing the cryptic epitope R39-R43 (purple) to the solvent. The O-circular form features an intramolecular interaction between DI (blue) and DV (yellow) with amino acids K19, R39 and R43 in DI potentially making contact with K305 and K317 in DV. The S-twisted conformation features a rotation of the DI/DII module, brokered by Lnk2, relative to the rest of the protein resulting in the blockade of the immunogenic region R39-R43 by the N-linked glycosylation (orange line).

While significant efforts have been made over the past decade to identify the cellular targets and signaling pathways triggered by anti-β_2_GPI antibodies(6, 14–17), our knowledge of the structural properties of β_2_GPI and the structural determinants for antigen-antibody recognition has lagged behind. Only two major advances came in recent years. First, it was discovered that anti-β_2_GPI antibodies in strong correlation with thrombosis recognize a conformational epitope comprising residues R39-R43 in the N-terminal domain, Domain I (DI), of β_2_GPI(18–22). Importantly this epitope was also proposed to be cryptic based on the evidence that anti-DI antibodies showed no reactivity against β_2_GPI in solution but did react well when β_2_GPI was immobilized onto hydrophilic surfaces or plastic plates pre-coated with negatively charged phospholipids(18, 23). Second, using X-ray crystallography(24, 25), small-angle X-ray scattering (SAXS)(26), negative stain electron microscopy (EM)(27), and atomic force microscopy (AFM)(28), it was found that β_2_GPI can adopt O-circular, S-twisted and J-elongated conformations according to its ligation status. (Fig. 1B). Given the poor reactivity of β_2_GPI toward anti-DI antibodies in solution as compared to β_2_GPI bound to negatively charged surfaces(19), the high salt conditions used in the crystallization experiments(24, 25) and the harsh purification method used to purify the protein used in the X-ray and SAXS studies, the O-circular form is currently regarded as the predominant conformation that the protein adopts under physiological pH and salt concentrations, which is immunologically inert, incapable of reacting against anti-DI antibodies(27). In contrast, the J-elongated form is believed to represent the immunogenic conformation of the protein, which appears when β_2_GPI is bound to the membranes. Dimeric complexes of the elongated form assembled on the activated endothelium(29) and stabilized by anti-β_2_GPI antibodies would then engage cellular receptors, such as Annexin A2, TLR4, ApoER2, and GpIb⍺, to induce a procoagulant and proinflammatory state(17, 30–34). Since the S-twisted conformation of the protein inferred by SAXS was detected neither by EM, nor by AFM, nor by X-ray crystallography, the current model also predicts that the S-twisted form represents a transient, unreactive intermediate state that the protein populates while transitioning between the J- and O-forms.

Although very popular in the APS field, it is important to acknowledge that the structural features of the O- and S-conformations are poorly defined due to the low-resolution models generated by EM, AFM and SAXS. Consequently, it remains unclear under what circumstances and how these forms interconvert, what is their physiological role, and how they participate in the mechanism of autoantibody recognition. Encouraged by our recent results with prothrombin(35–37), the second most common antigen of aPL in APS, this work was initiated to investigate the structural and conformational properties of β_2_GPI under conditions relevant to physiology and provide new insights into the mechanism of autoantibody recognition. Our results based on X-ray crystallography, single-molecule FRET (smFRET), SAXS, binding kinetics, and mutational studies unexpectedly reveal that human recombinant β_2_GPI adopts an elongated, flexible conformation, not circular, in which DI is exposed to the solvent and therefore available for autoantibody recognition. Based on this new evidence and previous findings, an alternative mechanism to explain how negatively charged phospholipids may enhance the affinity toward anti-DI autoantibodies without requiring opening of the protein structure or relocation of the glycosylations away from DI is proposed, and its implication to our understanding of APS discussed.

## Results

### Expression, purification, and functional characterization of human recombinant beta-2 glycoprotein I

Like beads on a string, CCP-domains are known to adopt a variety of orientations depending on the length of the linker connecting two adjacent domains and electrostatic properties(38). To get a better grasp of the structural architecture of β_2_GPI under conditions relevant to physiology, we set out to perform structural and biophysical studies of fully glycosylated human recombinant β_2_GPI. Two versions of the proteins were successfully expressed and purified under native conditions at high yield and purity. The first version, called LT-β_2_GPI, contained a long multifunctional cleavable tag at the N-terminus, located right before the natural N-terminal sequence ^1^GRTC^4^ (Fig. 2A). The tag was then cleaved with enterokinase to generate the intact, mature protein (hrβ_2_GPI). Removal of the tag was confirmed by N-terminal sequencing (Fig. 2B). The second version, called ST-β_2_GPI, contains a shorter, non-cleavable purification tag at the N-terminus that, based on our previous work(36), is expected not to affect the conformational properties of the protein (Fig. 2B). ST-β_2_GPI was made to eliminate the enterokinase cleavage step that was very laborious and not as efficient as expected. The presence of the short tag was confirmed by N-terminal sequencing and accounted for the different electrophoretic mobility observed between recombinant and plasma purified protein before and after enzymatic removal of the N-glycosylations (Fig. 2B).

**Figure 2.**
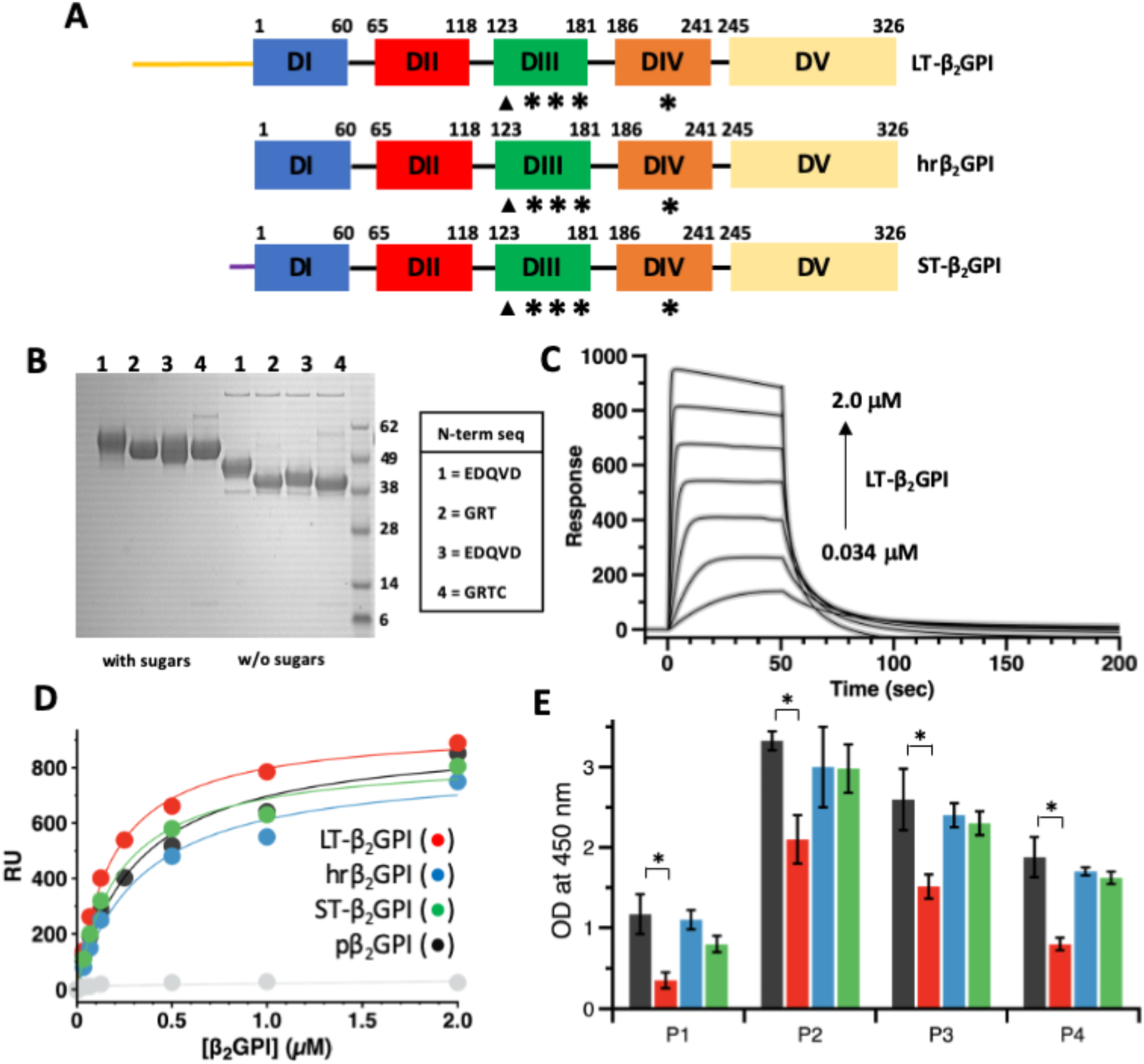
Functional characterization of human recombinant β_2_GPI. **(A)** Color-coded domain structure of the recombinant variants used in this work (i.e., LT-β_2_GPI, hrβ_2_GPI and ST-β_2_GPI) highlighting the position and chemical composition of the N-terminal tag. The long tag (LT, yellow) is composed of three parts: i) a calcium-dependent epitope for the monoclonal antibody HPC4 (EDQVDPRLIDGK); ii) a site-specific biotinylation sequence (AviTag); and iii) a conventional enterokinase recognition site (DDDDK). Two flexible linkers (i.e., GGGS) were introduced to separate the three functional units of the tag to avoid the formation of secondary structure and ensure exposure of the tag to solvent. Removal of the LT with enterokinase generates hrβ_2_GPI. The short tag version of β_2_GPI contains only the HPC4 purification tag (purple). **(B)** SDS-PAGE analysis of the recombinant proteins (sample 1=LT-β_2_GPI; sample 2=hrβ_2_GPI; sample 3=ST-β_2_GPI) and plasma purified β_2_GPI (sample 4) before (left) and after (right) the removal of the glycosylations. Protein Deglycosylation Mix II from NEB was used to remove O-linked and N-linked glycosylations under denaturing conditions. Chemical identify was verified by N-terminal sequencing and results are as follows: band 1=EDQVD; band 2=GRT; band 3=EDQVD; band 4=GRTC. **(C)** Representative sensograms of the interaction between LT-β_2_GPI and liposomes (PC:PS 80:20) monitored by SPR. Liposomes were immobilized on a L1 chip and soluble β_2_GPI (0-2 μM) was used in the fluid phase **(D)** Dose-dependent curves quantifying the interaction of LT-β_2_GPI (red circles), hrβ_2_GPI (blue circles) and ST-β_2_GPI (green circles) and pβ_2_GPI (gray circles) with liposomes monitored by SPR. Affinity values (K_d_) are 0.19±0.05 μM for LT-β_2_GPI, 0.33±0.08 μM for hrβ_2_GPI; 0.22±0.08 μM for ST-β_2_GPI and 0.32±0.09 μM for pβ_2_GPI. No significant binding was observed with liposomes entirely made of PC (light gray circles). Each experiment was repeated at least three times, using two distinct batches of proteins. **(E)** Reactivity of immobilized LT-β_2_GPI (red bars), hrβ_2_GPI (blue bars) and ST-β_2_GPI (green bars) and pβ_2_GPI (gray bars) against IgG anti-β_2_GPI antibodies followed by ELISA. Comparisons between 2 groups were performed using a Two-sample t-Test. Results were considered significant at p<.05 (*).

To evaluate the functional integrity of the recombinant proteins, LT-β_2_GPI, hrβ_2_GPI and ST-β_2_GPI were tested side by side with plasma purified β_2_GPI (pβ_2_GPI) in several biochemical assays. Using surface plasmon resonance (SPR), we found that all variants interacted avidly with liposomes containing negatively charged phospholipids such as phosphatidylserine, yet they failed to interact with phospholipids entirely made of phosphatidylcholine (Fig. 2C-D). Importantly, the values of the affinity constants were similar for all the constructs and consistent with published data(39), and so was the inhibitory effect of physiological concentrations of calcium chloride. These results document structural integrity of the hydrophobic loop in DV and also prove that the phospholipid binding activity of β_2_GPI is not perturbed by the presence or removal of the purification tags.

In addition to properly interacting with phospholipids, the recombinant proteins were also successfully recognized in ELISA assays by aPL isolated from four triple positive APS patients, which contain anti-DI antibodies(22, 35, 40) (Fig. 2E). In this case, however, LT-β_2_GPI exhibited significantly lower values of OD_450_ nm compared to the other variants and plasma purified protein, suggesting that the presence of the long tag may mask some epitopes or, more likely, change the preferential orientation of the molecule that is adsorbed onto the plastic surface. Taken together, these studies validate recombinantly made β_2_GPI as a proxy for plasma-purified β_2_GPI. They also demonstrate that, under physiological conditions, β_2_GPI is primed for phospholipid and heparin binding.

### X-ray crystal structure of human recombinant beta-2 glycoprotein I

To investigate the structural properties of the recombinant proteins, crystallization experiments were performed for all the protein constructs. While it was not possible to crystallize LT-β_2_GPI, we solved the X-ray crystal structures of hrβ_2_GPI, ST-β_2_GPI and pβ_2_GPI at 2.6, 3.0 and 2.4 Å resolution, respectively (Fig. 3A-C). Diffraction quality crystals were obtained after two weeks at 4°C using ammonium sulfate as a precipitating agent. The crystals belong to the orthorhombic space group C222_1_ (Table 1). Notably, ST-β_2_GPI, for which extra electron density was observed at the N-terminus (Fig. 3D), crystallized under similar conditions and in the same space group compared to hrβ_2_GPI and pβ_2_GPI, confirming minimal structural perturbation introduced by the artificial tag. β_2_GPI contains 22 cysteine residues. In our structures, regardless of the biological source and method of purification, all of them are engaged in 11 disulfide bridges (Fig. 3A-C). Given that purification of the recombinant proteins occurs under native conditions, this result indicates that, right after cell secretion, β_2_GPI does not contain free thiols.

**Table 1.**
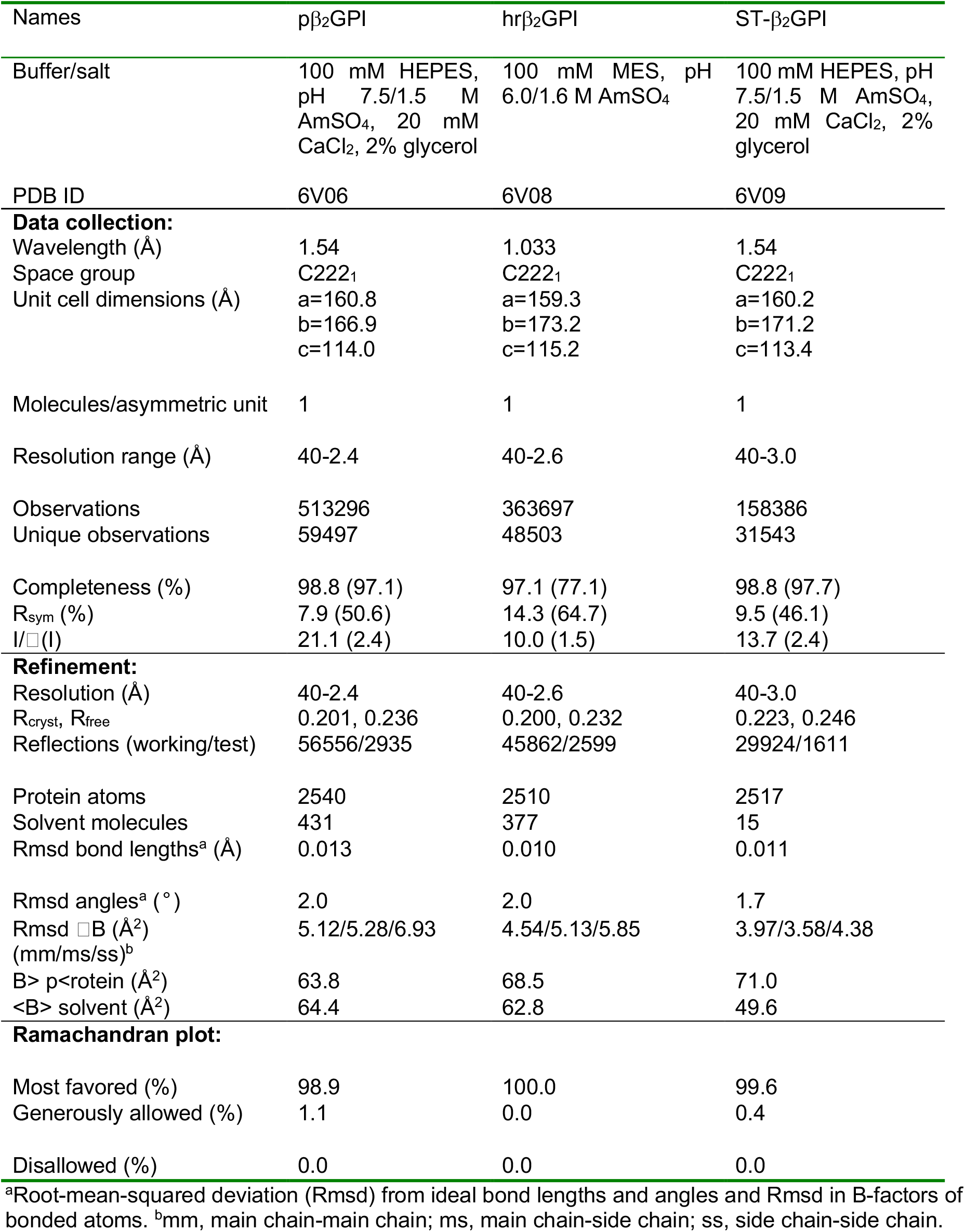
Crystallographic data for the structures of human beta-2 glycoprotein I

**Figure 3.**
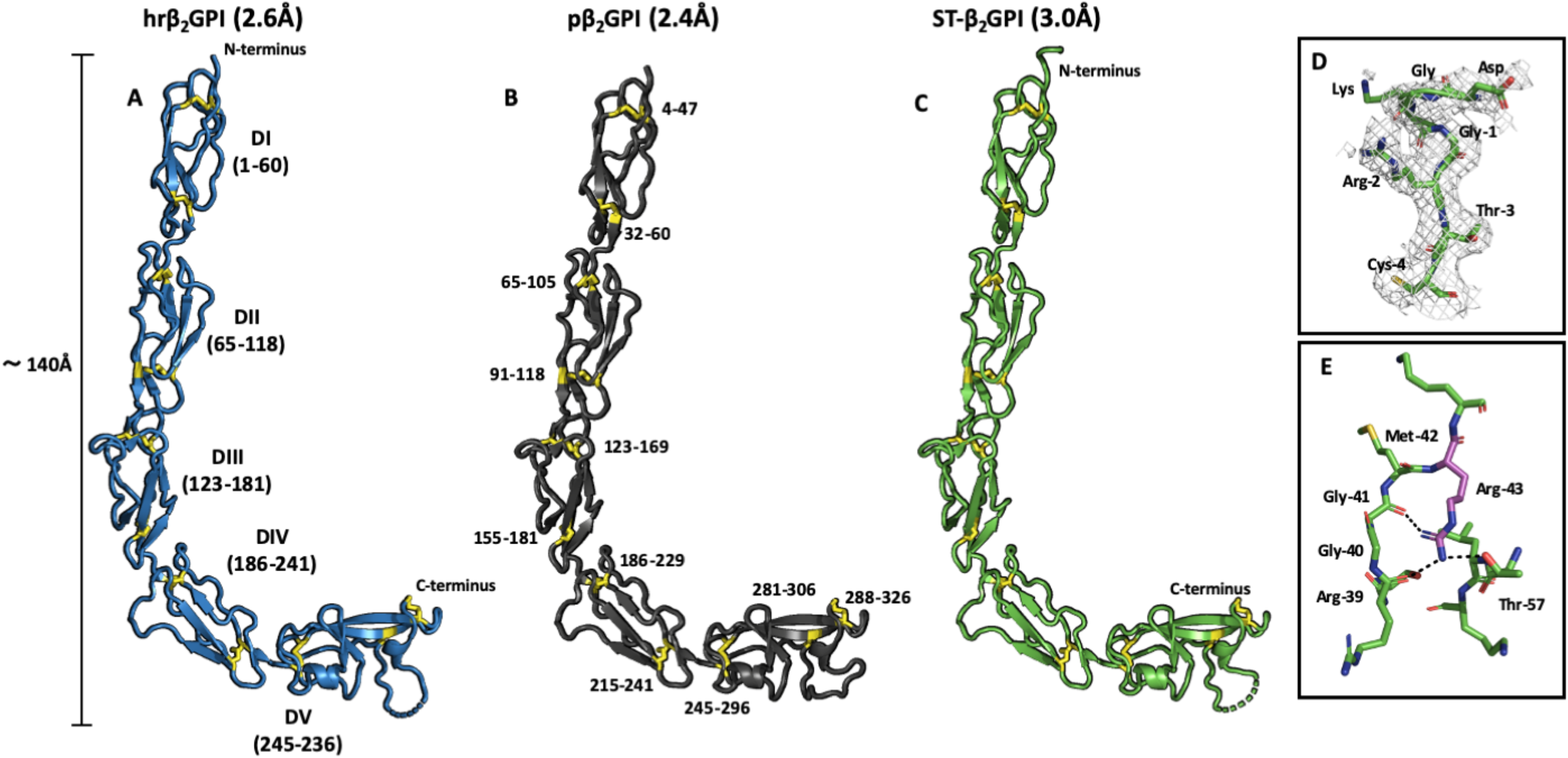
X-ray crystal structures of human recombinant β_2_GPI. X-ray crystal structures of **(A)** hrβ_2_GPI (blue), **(B)** pβ_2_GPI (green) and **(C)** ST-β_2_GPI (green) solved at 2.6 Å, 3.0 and 2.4 Å resolution. All three structures document similar elongated conformations of the protein spanning ~140 Å in length. Disulfide bonds are highlighted in yellow. **(D)** Zoom-in of three extra residues (DGK) belonging to the N-terminal tag preceding the natural sequence of β_2_GPI (^1^GRTC^4^) were exclusively found in the structure of ST-β_2_GPI, as expected. The electron density 2F_0_-Fc map is countered at 1.0σ. **(E)** Structural architecture of the epitope R39-R43 in DI highlighting the position and interactions of R43 (magenta stick) with the nearby residues R39, G41 and T57. Hydrogen bonds between the guanidinium group of R43 and neighboring residues are shown in black. Of note, the conformation of this segment is not involved in crystal contacts and is conserved in all the available crystal structures of β_2_GPI solved thus far, despite the high salt concentrations in which the crystals grow, suggesting that this is a genuine structural feature of β_2_-GPI.

All three independently solved X-ray crystal structures depicted β_2_GPI featuring an elongated conformation spanning ~140 Å in length, from the N- to the C-terminus. The first three domains, DI-DIII, are aligned along the vertical axis of the molecule, whereas DIV and DV bend, forcing the molecule to adopt a characteristic J-shaped elongated form resembling a hockey stick. DI and DV are located >100 Å apart and both of them are exposed to the solvent. Interestingly, however, the side chain of residue R43, which is part of the cryptic epitope recognized by anti-DI antibodies, is not exposed to the solvent and is part of a hydrogen bond network made up by residues R39, G41 and T57 (Fig. 3E).

Overall, the three new structures are superimposable and similar to the published ones(24, 25), yet they are not identical (Fig. 4A). One significant difference regards the conformation the hydrophobic loop in DV (residues 308-319), which, given its flexibility and exposure to the solvent, varies in every structure. Another difference is the significant extra electron density after molecular replacement in the datasets at a higher resolution (i.e., 2.4 and 2.6Å), suggesting that the input structural model used to solve the structures (1C1Z(25)) was incomplete (Fig. 4B). We attributed this density to the N-linked glycosylations (Fig. 4C). Modeling of the glycans provided a more complete view of the glycoprotein and offered new important clues on how β_2_GPI may adopt multiple conformations in solution and become immunogenic. Our new structural model predicts that epitopes in DII, DIII and DIV are mostly buried by the presence of the glycosylations, whereas DI and DV are not. It also suggests that a conformational change may be required to expose R43 to the solvent. Due to the presence of sialic acid, N-glycosylations typically are negatively charged. DI and DV, in contrast, are positively charged (Fig. 4D). It is therefore reasonable to admit that the presence of glycosylations affects the conformational landscape of the molecule 1) by limiting the number of spatial arrangements that the protein can adopt in solution because of steric hindrance, 2) by facilitating looping of the molecule by neutralizing the repulsive positive electrostatic potential of DI and DV and 3) together with DIV, by preserving an elongated conformation of the protein when the protein is anchored to the lipids via DV.

**Figure 4.**
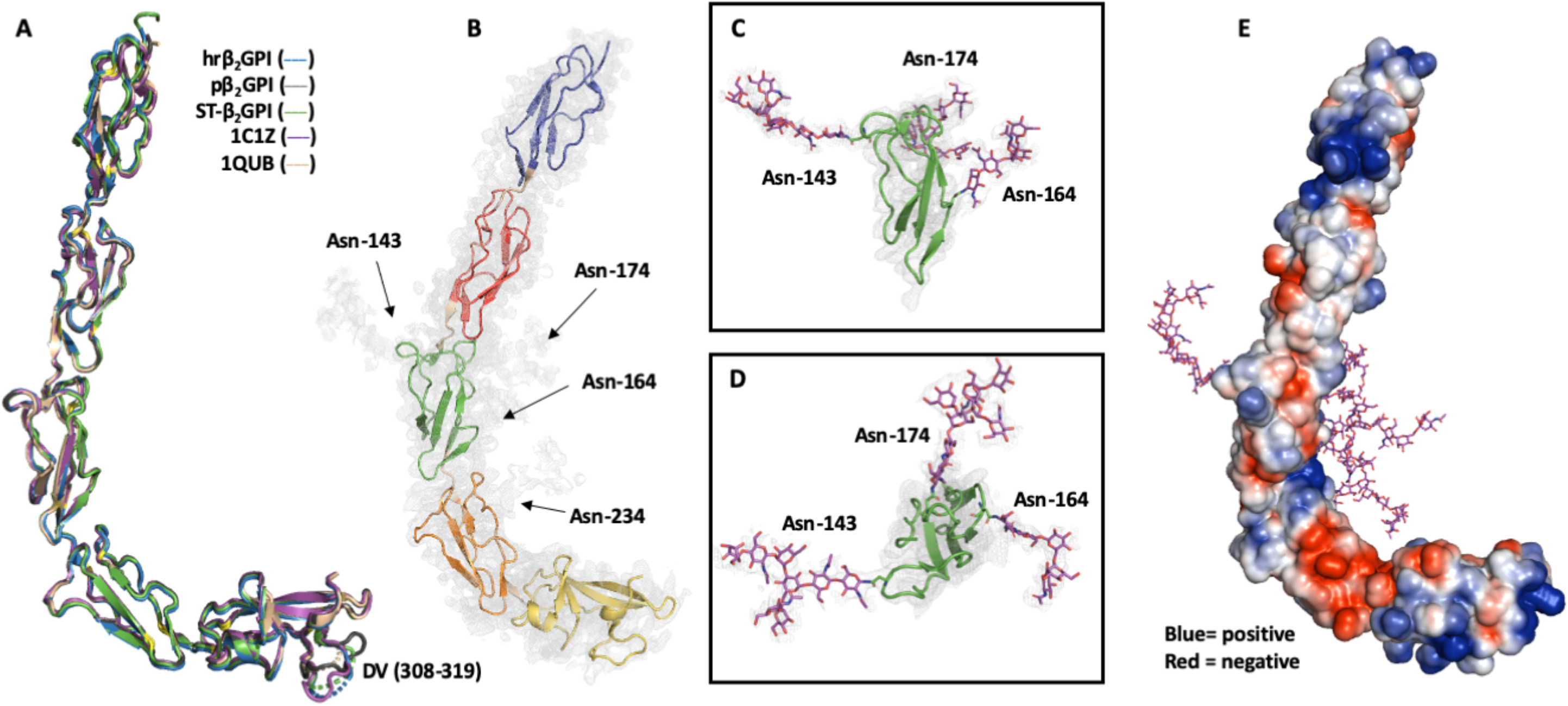
Location and structural role of the N-linked glycosylations. **(A)** Superposition of five X-ray crystal structures of β_2_GPI highlights structural similarities yet diversity of the phospholipid binding loop in DV (residues 308-319). **(B)** Extra electron density detected in the structure of pβ_2_GPI solved at 2.4Å resolution attributed to the N-linked glycosylations. The domains of β_2_GPI are color coded as shown in Figure 1. Guided by mass spectrometry analyses(66), we modeled the following sugar sequences: Gal_2_GlcNAc_2_Man_3_GlcNAc_2_ at N143, Gal_2_GlcNAc_2_Man_3_GlcNAc_2_ at N164, GlcNAc_2_ at N174 and Gal_3_GlcNAc_2_ at N234. The presence of a putative O-linked glycosylation at T130 could not be confirmed because of weak density. **(C)** Side and top **(D)** views of the N-glycosylations surrounding DIII. The electron density 2F_0_-Fc map is countered at 0.8σ. **(E)** Asymmetric distribution of the electrostatic potential displaying positive (blue) and negative (red) clusters with a −2.0-2.0-intensity scale. The N-glycosylations are shown as magenta sticks.

### Solution structure of human recombinant beta-2 glycoprotein I

It is generally believed that the J-elongated conformation trapped in the X-ray studies is not a genuine representation of the protein structure in solution(2, 26). The high ionic strength used in the crystallization buffers may destabilize hydrogen bonds and favor hydrophobic interactions thus forcing the protein to assume a non-native conformation(2). To address this concern, we applied smFRET to β_2_GPI(37, 41, 42). By recording the energy that is transferred from an excited molecule (Donor) to a second molecule with spectral overlap (Acceptor) at the single molecule level, smFRET measures distances on a nanometer scale thus serving as a molecular ruler.

Guided by our new structures, we generated four FRET pairs in the ST background, S13C/S112C, S13C/S312C, S112C/S312C and S190C/S312C (Fig. 5A-B), by substituting the natural serine residues with isosteric cysteines and then reacting those newly engineered cysteines with Alexa Fluor 555 (AF555) maleimide as FRET donor and Alexa Fluor 647 (AF647) maleimide as FRET acceptor. Residue 13 is located in DI, residue 112 is located in DII, residue 190 is located in DIV and residue 312 is located in DV. Labeling occurred only at the engineered sites as no fluorescence was observed for β2GPI wild type (Fig. 5C). This result is consistent with our structural data and previous findings(43) documenting that β_2_GPI wild type does not contain free thiols. Given the Förster radius R0 = 50 Å of the 555/647 FRET couple, the crystal structure predicts no FRET for the mutants S112C/S312C and S13C/S312C. In contrast, high FRET and low but measurable FRET values are expected for the FRET pair S13C/S112C and S190C/S312C, respectively. This is because residues 13 and 112 are located ~24 Å apart while the Cα-Cα distance between residues 190 and 312 is ~46Å. Remarkably, the experimental results were fully consisted with our structure-based calculations, thus validating the elongated conformation in solution and unequivocally proving that this conformations predominates (>90%) under physiological pH and salt concentrations (Fig. 5D). Specifically, probes located at positions C13 and C312 and C112 and C312 reported a negligible FRET signal, whereas probes attached to the S13C/S112C and S190C/S312C mutants reported high (E_FRET_=0.92) and low (E_FRET_=0.26) FRET values, respectively. Interestingly, the construct 190/312 displayed a FRET distribution wider than the theoretical distribution predicted by shot noise(44) documenting the existence of multiple conformations at equilibrium brokered by the flexibility of Lnk4. Importantly, no significant FRET differences were observed neither in the presence of high (1.5 M) or low (25 mM) concentrations of sodium chloride nor under acidic (pH 3.4) or alkaline (pH 11.0) pH, suggesting that, in contrast to previous findings, variation of the ionic strength and pH produces minor conformational changes that could not be detected by our FRET pairs.

**Figure 5.**
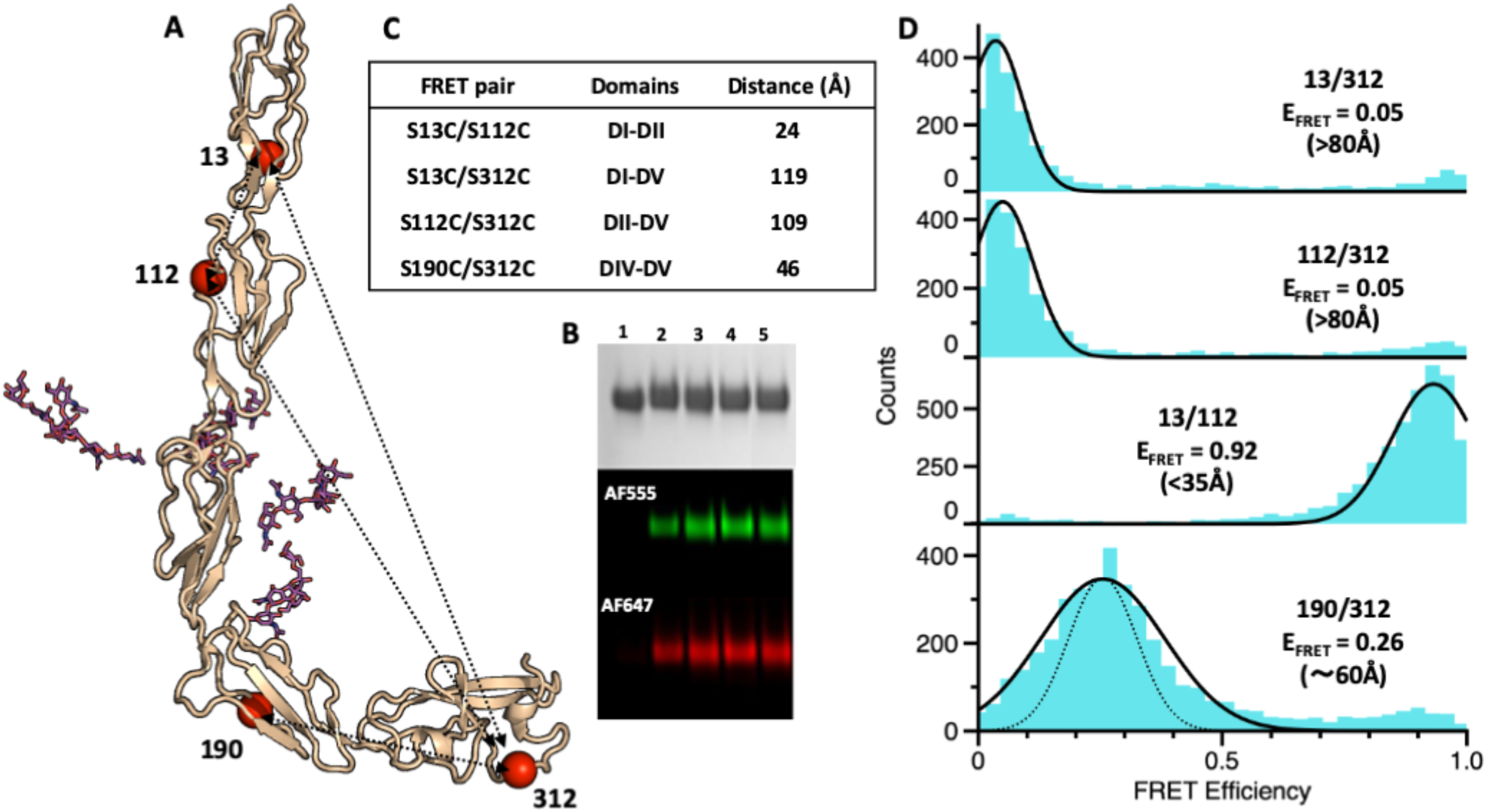
smFRET measurements of β_2_GPI in solution. **(A)** Structure-based design of the FRET constructs S13C/S112C, S13C/S312C, S112C/S312C, and S190C/S312C used in this study. Ser residues mutated to Cys for conjugation with the thiol-reactive dyes AF555 and AF647 used in smFRET measurements are indicated by red spheres. **(B)** FRET couples are listed with their respective domains. Cα-Cα distances obtained from the crystal structure of hrβ_2_GPI. **(C)** After labeling and gel filtration, selective incorporation of the fluorescence dyes was verified by loading the proteins (samples 2-5) alongside β_2_GPI WT (sample 1), into a gradient 4 –12% polyacrylamide gel in the presence of SDS and visualized by Coomassie Brilliant Blue R-250 (black and white) or fluorescence intensity by exciting donor at 532 nm (red panel) and acceptor at 640 nm (blue panel). **(C)** smFRET histograms of the mutants S13C/S312C, S112C/S312C, S13C/S112C, and S190C/S312C labeled with AF555/647 measured in Tris 20 mM pH 7.4, 145 mM NaCl, 0.1% Tween 20 for 1 hour at room temperature at a concentration of 100 pM. Populations were fit to a single Gaussian distribution (black lines). FRET efficiency values and calculated distances are indicated. The theoretical shot noise peak highlighting conformational heterogeneity for the FRET pair 190/312 is shown as dotted line.

To rule out potential artifacts arising from the substitution of natural amino acids with cysteine and incorporation of fluorescent dyes, we collected SAXS data for the ST-β_2_GPI and pβ_2_GPI, under physiological conditions (Fig. 6A-B). SAXS is a biophysical method that is particularly useful to assess the overall shape of biological macromolecules in solution(45), i.e., linear vs. globular, and, similar to smFRET, is therefore ideal to detect large conformational changes in β_2_GPI. Both the radius of gyration (Rg) and the computed pair distance distribution functions p(r) were fully consistent with interpretation provided by the smFRET experiments and previous SAXS measurements (26) suggesting an extended, flexible protein conformation in solution. Furthermore, the scattering profile of the recombinant and plasma proteins were very similar, confirming structural equivalency between the two proteins.

**Figure 6.**
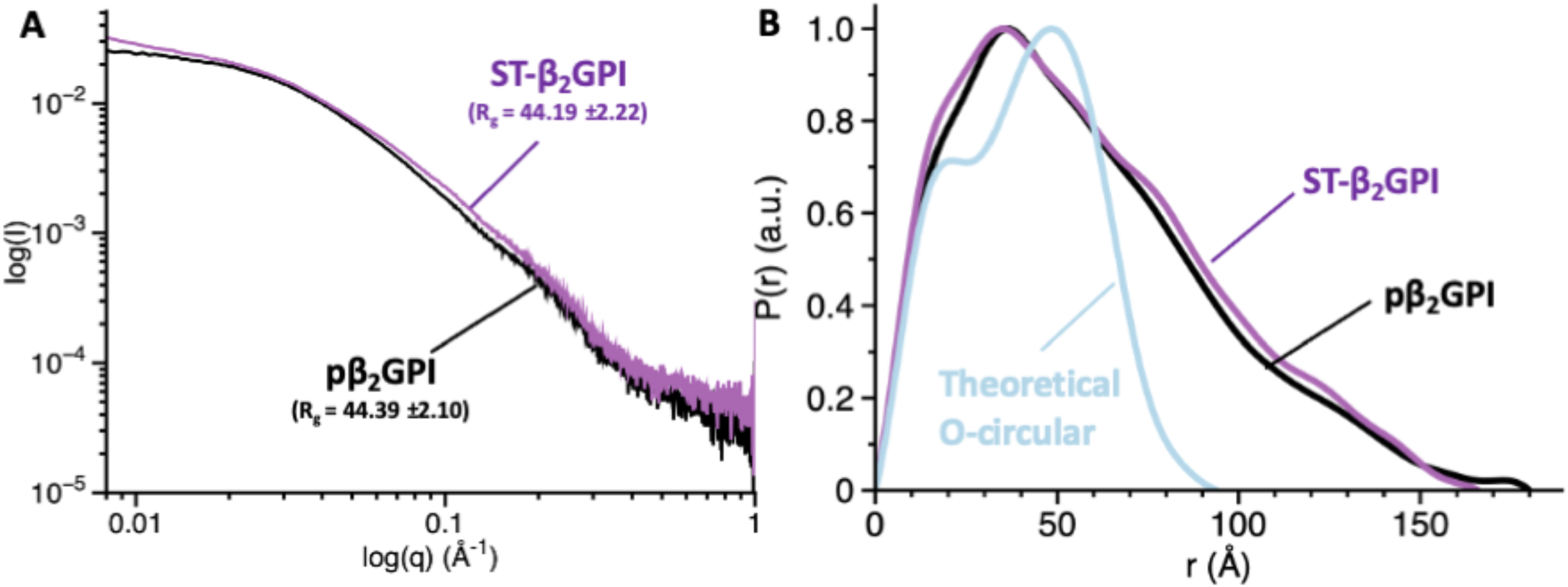
Elongated conformation of β_2_GPI revealed by SAXS. Scattering profiles **(A)** and pair distribution functions **(B)** for pβ_2_GPI (black) and ST-β_2_GPI (magenta) collected at 2 mg/ml under physiological conditions (Tris 20 mM pH 7.4, 145 mM NaCl). The calculated values of the radius of gyration, Rg, are very similar for pβ_2_GPI and ST-β_2_GPI. The blue curve in panel B, which is significantly different from the experimental scattering profiles, represents the theoretical pair distribution function for a hypothetical circular conformation.

### Autoantibody binding studies

In addition to demonstrating that β_2_GPI adopts an elongated conformation in solution, our structural analysis predicts that this form may be primed for autoantibodies binding, especially anti-DI antibodies. To test this hypothesis, we took advantage of the reactivity of MBB2(13), a newly developed recombinant monoclonal antibody raised against DI that, upon complement fixation, recapitulates, *in vivo*, most of the clinical characteristics assigned to pathogenic aPL. The binding of β_2_GPI to MBB2 was followed using SPR (Fig. 7), a technique that allows to measure association (on) and dissociation (off) rate constants in real-time thus enabling a deeper understanding of the chemical nature of the molecular interaction.

**Figure 7.**
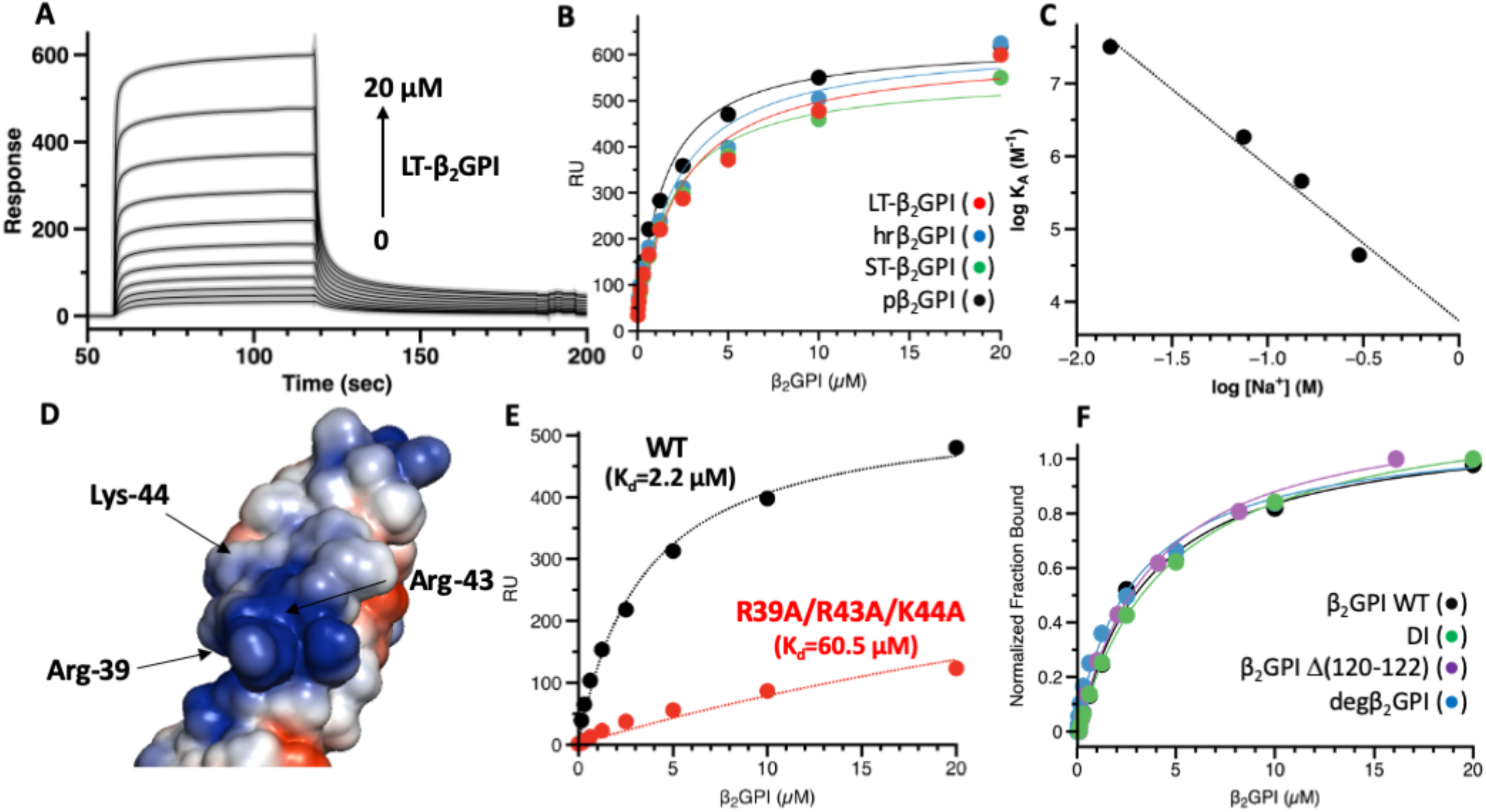
Exposure of DI revealed by MBB2 binding and mutagenesis studies. **(A)** Interaction of MBB2 and LT-β_2_GPI monitored by SPR. MBB2 was immobilized onto a CM5 chip using NHS/EDC chemistry to a final density of 6000 RU. A solution of LT-β_2_GPI (0-20 μM) in Tris 20 mM pH 7.4, 145 mM NaCl 0.01% Tween 20 was injected at 30 μl/min for 60 sec to observe binding followed by 60 sec dissociation in running buffer. **(B)** Dose-dependent curves quantifying the interaction of LT-β_2_GPI (red circles), hrβ_2_GPI (blue circles) and ST-β_2_GPI (green circles) and pβ_2_GPI (gray circles) with MBB2 monitored by SPR. Affinity values (K_d_) are 2.2±0.5 μM for LT-β_2_GPI, 2.1±0.6 μM for hrβ_2_GPI; 1.8 ±0.5 μM for ST-β_2_GPI and 1.4±0.6 μM for pβ_2_GPI. Each experiment was repeated at least three times, using two distinct batches of proteins. **(C)** Effect of the ionic strength. SPR binding experiments were performed at 15, 75, 150 and 300 mM NaCl. Analysis of the slope of the linear fit of the association constant vs Na+ reveals a strong dependency of the ionic strength and the presence of at least 2.0 ionic contacts (salt bridges) in the complex(66, 67). **(D)** Location of residues R39, R43 and K44 targeted by site-directed mutagenesis. **(E)** SPR analysis of β_2_GPI WT and mutant R39A/R43A/RK44A reveal that residues R39 and R43 are critical for MBB2 binding. **(F)** SPR analysis of β_2_GPI WT (black circles), DI (green circles), β_2_GPIΔ(120-122) (magenta circles) and degβ_2_GPI (blue circles). Removal of DII-DV, perturbation of Lnk2 and removal of the glycosylations do not affect the binding affinity of MBB2 towards β_2_GPI. This indicates that 1) DI is primed for autoantibody recognition in solution 2) the exposure of residues R39 and R43 in DI is independent of the conformation of Lnk2 and 3) Lnk2 is not an epitope of MBB2. Given the different molecular weight between the constructs, the binding curves are reported as normalized fraction bound vs analyte to facilitate comparison. Each experiment was repeated at least three times, using two distinct batches of proteins.

To retain the native conformation of β_2_GPI in solution, we immobilized MBB2 to the chip’s surface and injected β_2_GPI in the fluid phase. Binding between MBB2 and β_2_GPI should occur only if DI is exposed to the solvent. This experimental setup is different from previously reported interaction data between MBB2 and β_2_GPI in which β_2_GPI was covalently immobilized on a dextran-based chip and the antibody was used in the fluid-phase to mimic binding of MBB2 to β_2_GPI bound to negatively charged phospholipids(13). The results of the experiments shown in Fig. 7A demonstrate that MBB2 interacts with β_2_GPI in solution with a modest but measurable affinity, characterized by a dissociation constant K_d_=2.2±0.2 μM (Fig. 7B). Remarkably, the value of K_d_ determined for MBB2 is similar to the value of K_d_ obtained by Dienava-Verdoold et al. for patient-derived monoclonal antibodies targeting Domain I (46), yet it is 200-fold weaker than the affinity previously determined for MBB2 towards immobilized β_2_GPI (i.e., Kd=11 nM)(13). Such as dramatic difference is expected for aPL found in APS patients, which prefer immobilized β_2_GPI, thus making MBB2 is a highly relevant tool for biochemical investigations.

Interestingly, the interaction between MBB2 and soluble β_2_GPI was characterized by very fast on and off rates, suggesting that electrostatic interactions may dominate the binding interface. This was demonstrated by systematic experiments in which we varied the ionic strength of the running buffer from 300 to 15 mM NaCl. As expected, the affinity constant of MBB2 for β_2_GPI (K_A_) increased ~700-fold at low salt concentrations, from 4.4*10^4^ M^−1^ at 300 mM NaCl to 3.2*10^7^ M^−1^ at 15 mM NaCl (Fig. 7C). Importantly, the higher affinity of MBBS for β_2_GPI at low salt originated from a substantial reduction of the off-rate whereas the on-rate remained mostly unaffected. A significant ~30-fold reduction of the affinity under physiological conditions was also detected after mutating the positively charged residues R39, R43 and K44 in DI with the neutral amino acid alanine (Fig. 7D-E), confirming the electrostatic nature of such interaction and the ability of MBB2 to interact with an epitope of DI that is targeted by pathogenic aPL.

Previous studies have proposed that the epitope R39-R43 is cryptic because it is buried by DV in the O-circular form(27) or by the N-linked glycosylations in the S-twisted form(26). To test these hypotheses, we measured the affinity of three new constructs, isolated DI (residues 1-60), β_2_GPI deletion linker 2 (β_2_GPI Δ120-122) and recombinantly deglycosylated β_2_GPI (degβ_2_GPI, T130S/N143Q/N164Q/N174Q/N234Q/) toward immobilized MBB2. Our binding data shown in Fig. 7F indicates that all three constructs interact with immobilized MBB2 with micromolar affinity comparable to full-length β_2_GPI wild-type thus ruling out a significant contribution of the neighboring domains and the glycosylations in shielding the R39-R43 epitope when the protein is free in solution. These constructs were indeed designed based on previous knowledge to force the exposure of DI to the solvent and, in principle, should have had higher affinity for MBB2.

## Discussion

Owing to its flexibility, the structural architecture of β_2_GPI has remained controversial, and so is the mechanism of autoantibody recognition. A first major conclusion emerging from this study is that human recombinant β_2_GPI expressed in mammalian cells and purified under native conditions adopts an elongated, flexible conformation in which DV and DI are exposed to the solvent. In the free form, under physiological pH and salt concentrations, β_2_GPI is therefore primed for phospholipid binding and autoantibody recognition. A second major conclusion is that the recombinant protein is structurally and functionally identical to β_2_GPI purified from plasma using the perchloric acid method. Hence, the elongated conformation of β_2_GPI is not an artifact caused by the harsh purification methods or crystallization conditions but a genuine conformation of the protein in solution.

The recognition that the elongated form of β_2_GPI exists and, according to our smFRET experiments, perhaps predominates in human plasma bears important implications in our understanding of the APS pathology. It also provides new ideas for the development of APS-focused diagnostics and therapeutics. Regarding the mechanism of anti-DI antibody recognition, our structural and binding data indicates that opening of the protein structure and relocation of DI away from the glycosylations are neither necessary nor sufficient to explain how β_2_GPI becomes a better antigen for anti-DI antibodies upon binding to the membranes. They instead strongly suggest that, in agreement with previous models(39, 47, 48), binding of the pre-existing elongated conformation of β_2_GPI to the membranes gives rise to an ideal surface in which β_2_GPI has a sufficiently high density and adopts a favorable orientation that promotes bivalent binding. In this context, rotation or bending of the CCP domains relative to the plane of the membrane such as those documented here by the FRET couple 190/312 may be important for proper packing of β_2_GPI onto the lipids and, in agreement with previous studies(23, 49), they may even promote oligomerization. A contribution of local conformational changes, such as exposure of R43 upon binding to the lipids, is also possible, yet, considering the intrinsic low affinity of such autoantibodies for their targets, the modest effect caused by mutations in isolated DI(50), and the key role of bivalency documented before(48, 51), the energetic contribution of this process is expected to be minor. The fact that immunocomplexes are very difficult to detect in patients’ plasma(52) even though β_2_GPI is primed for autoantibody binding, is a consequence of the low affinity, fast binding kinetics and low abundance of such autoantibodies in APS patients (0.5% of the total IgG or less). Since the low affinity arises from a very fast dissociation rate constant, the immunogenic complexes are unstable in the fluid phase as they dissociate very rapidly. In this context, a possible role of the negatively charged surfaces suggested by our SPR binding experiments performed at low ionic strength would be to stimulate the binding of anti-DI antibodies to β_2_GPI by slowing down the dissociation rate, thus resulting in complex stabilization. Such a mechanism is fully consistent with recent and previous data(35, 48), and might be conserved among other aPL.

Another important aspect of the pathogenesis of APS is the interaction of β_2_GPI/aPL complexes with cell receptors, mainly via DV, resulting in amplification of pro-inflammatory and pro-thrombotic responses(6, 17, 30, 33, 53, 54). Our structural model predicts that interaction of β_2_GPI with cell receptors may occur without the need of phospholipids, yet the signaling cascade may be triggered by receptor dimerization, which is induced by aPL(17, 33, 55). The need for β_2_GPI, aPL, and suitable receptors explain why the clustering of β_2_GPI onto negatively charged phospholipids is not sufficient to trigger cell activation(29) and why binding of β_2_GPI to cell receptors and anionic phospholipids is mutually exclusive(54).

Regarding the development of new diagnostics and therapeutics, if the main role of the membranes is to increase the local concentration of the elongated form, we speculate that immobilization of human recombinant β_2_GPI produced in this work at the desired density and with a defined orientation should provide a novel, efficient, robust and cost-effective method to detect anti-β_2_GPI antibodies. On the other side, blocking the binding of the elongated form of β_2_GPI to cell receptors and phospholipids should limit the pathogenic effects of anti-DI antibodies. This approach could complement current strategies aimed at competing with anti-DI antibodies in solution(13, 56–58) as it would theoretically block other potentially pathogenic aPL, in addition to those targeting DI. Consistent with this premise, antibodies against DV found in APS patients do not induce thrombosis but are protective instead (59), and a novel dimeric molecule, A1-A1, protects mice from aPL-induced thrombosis by interfering with ApoER2 and phospholipid binding(60).

It is important to acknowledge that, even though all the structural and biochemical experiments were performed using highly purified proteins solubilized in physiological buffers, β_2_GPI has never been exposed to endothelial or circulating blood cells. Hence, it remains possible, although unlikely, that the O-circular form of β_2_GPI previously documented by EM and AFM studies but not detected by our structural studies, unless the result of an experimental artifact, may arise from chemical and/or posttranslational modifications catalyzed by membrane-bound proteins. Future studies will be needed to clarify this matter.

## Materials and Methods

### Protein production and purification

β_2_GPI wild-type and mutants were expressed in BHK and HEK293 mammalian cells and purified to homogeneity by immunopurification, heparin and size exclusion chromatography (SEC) after swapping the signal peptide of β_2_GPI with the one of the coagulation factor X to boost expression and adding a furin specific recognition motif RRKR for quantitative post-translational processing. The purity and chemical identity of each fragment were verified by SDS-PAGE and N-terminal sequencing. Domain I (1–60) was chemically synthesized and refolded as done before(57). Plasma-derived β_2_GPI (pβ_2_GPI) was purified using the perchloric acid method, as described previously(57). MBB2 was produced as described before(13). Liposomes composed of phosphatidylcholine (PC) or phosphatidylcholine and phosphatidylserine (PS) in a 4:1 molar ratio (PC:PS) were prepared by extrusion using 100 nm polycarbonate membranes (Avanti Polar Lipids, Alabaster, AL), kept a 4°C and used within 7 days. ELISA assays were performed as described before(22, 35, 57). Protein concentrations were determined by reading at 280 nm with molar extinction coefficients adjusted according to the amino acid sequence. All other chemicals were purchased from Sigma-Aldrich.

### Surface Plasmon Resonance (SPR) experiments

Binding affinities for liposomes were measured as done before(35) using L1 sensor chip in which liposomes were immobilized at 1600 RU. Titrations were performed by injecting increasing concentrations (0-2 μM) of β_2_GPI and its variants in running buffer (20 mM Tris pH 7.4, 150 mM NaCl, 0.1% w/w BSA) at a flow rate of 25 μl/min at 25°C. Binding affinities for MBB2 were measured using CM5 sensor chip in which MBB2 was immobilized at 6000 RU using NHS/EDC chemistry. Titrations were performed by injecting increasing concentrations (0-20 μM) of β_2_GPI and its variants in running buffer (20 mM Tris pH 7.4, 25-300 mM NaCl, 0.01% w/w Tween20) at a flow rate of 25 μl/min at 25°C. All experiments were carried out using on a BIAcore-S200 instrument (GE-Healthcare). The dissociation constants (Kd) were obtained as a fitting parameter by plotting the value of the response units (RU) at the steady state for each concentration using the BIAevaluation software and Origin Pro 2015.

### X-ray studies

Crystallization of human recombinant (hr β_2_GPI and ST-β_2_GPI) and pβ_2_GPI was achieved at 4°C by the vapor diffusion technique, using the Art Robbins Instruments PhoenixTM liquid handing robot with 10 mg/ml protein 0.3 μl mixed with an equal volume reservoir solution. Optimization of crystal growth was achieved by the hanging drop vapor diffusion method mixing 3 μl of protein (10 mg/ml) with equal volumes of reservoir solution at 4°C. After 7-10 days at 4°C, crystals were frozen with 25% glycerol from original mother liquid. X-ray diffraction data were collected at 100° K with a home source (Rigaku 1.2 kw MMX007 generator with VHF optics) Rigaku Raxis IV++ detector for pβ_2_GPI and ST-β_2_GPI, and with detector Pilatus of Beamline IDD23, at the Advanced Photon Source, Argonne, IL for hrβ_2_GPI. Datasets were indexed, integrated and scaled with the HKL2000 software package(61). All structures were solved by molecular replacement using PHASER from the CCP4 suite(62) and the structure of pβ_2_GPI (PDB ID code 1C1Z) as starting model. Refinement and electron density generation were performed with REFMAC5 from CCP4 package. 5% of the reflections were randomly selected as a test set for cross-validation for four structures. Model building and analysis of the structures were carried out using COOT(63). Ramachandran plots were calculated using PROCHECK. Statistics for data collection and refinement are summarized in Table 1. Atomic coordinates and structure factors have been deposited in Protein Data Bank (accession codes: 6V06 for pβ_2_GPI at 2.4Å, 6V08 for hrβ_2_GPI at 2.6Å, and 6V09 for ST-β_2_GPI at 3.0Å).

### Single-molecule FRET

Selective labeling of the unpaired Cys residues with Alexa Fluor 555-C2-maleimide as the donor and Alexa Fluor 647-C2-maleimide as the acceptor was achieved as described recently for prothrombin(36, 37). FRET measurements of freely diffusing single molecules were performed with a confocal microscope MicroTime 200 (PicoQuant, Berlin, Germany), as detailed elsewhere(36, 37).

### Small Angle X-ray Scattering Measurements

SAXS data were collected at the beamline 12-ID-B of the Advanced Photon Source at Argonne National Laboratory (Argonne, IL) on ST-β_2_GPI and pβ_2_GPI at concentrations of 0.5, 1, 2, and 5 mg/ml. The radius of gyration, Rg, was determined using the Guinier approximation in the low q region (qRg<1.3), and its linearity served as an initial assessment of data and sample quality. Maximum particle dimension, D_max_, and distance distribution function, P(r), were calculated using GNOM.

## Acknowledgments

This work was supported in part by the National Institutes of Health Research Grant HL150146 (N.P.) and President’s Research Fund, Saint Louis University (N.P.). This research used resources of the Advanced Photon Source, a U.S. Department of Energy (DOE) Office of Science User Facility operated for the DOE Office of Science by Argonne National Laboratory under Contract No. DE-AC02-06CH11357.

## Authorship Contributions

E.R., Z.C., and N.P. solved the X-ray crystal structures; E.R., W.P., M.C., and N.P. performed SPR experiments; M.C., W.P., E.R., and N.P. performed smFRET experiments; X.Z. and E.R. performed SAXS experiments; V.D.F. produced the isolated DI; P.M. and F.T produced the monoclonal antibody MBB2; V.P. provided plasma samples; E.R., Z.C., W.P., M.C., X.Z., R.K.A., K.R.M., F.T., and N.P critically analyzed the data; N.P. designed the research and wrote the manuscript. All authors reviewed the manuscript.

## References

1. H. E. Schultze, K. Heide, H. Haupt, Uber ein bischer unb ekanntesniedermolekularis b2-globulins des human serums.. Naturwissens-chaften 48, 719 (1961).

2. P. G. de Groot, J. C. Meijers, beta(2) -Glycoprotein I: evolution, structure and function. J Thromb Haemost 9, 1275–1284 (2011).

3. E. M. Bevers, M. Galli, Beta 2-glycoprotein I for binding of anticardiolipin antibodies to cardiolipin. Lancet 336, 952–953 (1990).

4. M. Galli et al., Anticardiolipin antibodies (ACA) directed not to cardiolipin but to a plasma protein cofactor. Lancet 335, 1544–1547 (1990).

5. H. P. McNeil, R. J. Simpson, C. N. Chesterman, S. A. Krilis, Anti-phospholipid antibodies are directed against a complex antigen that includes a lipid-binding inhibitor of coagulation: beta 2-glycoprotein I (apolipoprotein H). Proc Natl Acad Sci U S A 87, 4120–4124 (1990).

6. B. Giannakopoulos, S. A. Krilis, The pathogenesis of the antiphospholipid syndrome. N Engl J Med 368, 1033–1044 (2013).

7. J. S. Levine, D. W. Branch, J. Rauch, The antiphospholipid syndrome. N Engl J Med 346, 752–763 (2002).

8. T. Koike, E. Matsuura, Anticardiolipin antibodies and beta 2-glycoprotein I. Lupus 5, 156–157 (1996).

9. J. D. Oosting, R. H. Derksen, H. T. Entjes, B. N. Bouma, P. G. de Groot, Lupus anticoagulant activity is frequently dependent on the presence of beta 2-glycoprotein I. Thromb Haemost 67, 499–502 (1992).

10. F. Fischetti et al., Thrombus formation induced by antibodies to beta2-glycoprotein I is complement dependent and requires a priming factor. Blood 106, 2340–2346 (2005).

11. S. S. Pierangeli, E. N. Harris, In vivo models of thrombosis for the antiphospholipid syndrome. Lupus 5, 451–455 (1996).

12. A. Arad, V. Proulle, R. A. Furie, B. C. Furie, B. Furie, beta(2)-Glycoprotein-1 autoantibodies from patients with antiphospholipid syndrome are sufficient to potentiate arterial thrombus formation in a mouse model. Blood 117, 3453–3459 (2011).

13. C. Agostinis et al., A non-complement-fixing antibody to beta2 glycoprotein I as a novel therapy for antiphospholipid syndrome. Blood 123, 3478–3487 (2014).

14. N. Prinz et al., Antiphospholipid antibodies induce translocation of TLR7 and TLR8 to the endosome in human monocytes and plasmacytoid dendritic cells. Blood 118, 2322–2332 (2011).

15. N. Muller-Calleja et al., Complement C5 but not C3 is expendable for tissue factor activation by cofactor-independent antiphospholipid antibodies. Blood Adv 2, 979–986 (2018).

16. A. Hollerbach, N. Muller-Calleja, A. Canisius, C. Orning, K. J. Lackner, Induction of tissue factor expression by anti-beta2-glycoprotein I is mediated by tumor necrosis factor alpha. J Thromb Thrombolysis 49, 228–234 (2020).

17. K. L. Allen et al., A novel pathway for human endothelial cell activation by antiphospholipid/anti-beta2 glycoprotein I antibodies. Blood 119, 884–893 (2012).

18. B. de Laat, R. H. Derksen, R. T. Urbanus, P. G. de Groot, IgG antibodies that recognize epitope Gly40-Arg43 in domain I of beta 2-glycoprotein I cause LAC, and their presence correlates strongly with thrombosis. Blood 105, 1540–1545 (2005).

19. B. de Laat, R. H. Derksen, M. van Lummel, M. T. Pennings, P. G. de Groot, Pathogenic anti-beta2-glycoprotein I antibodies recognize domain I of beta2-glycoprotein I only after a conformational change. Blood 107, 1916–1924 (2006).

20. B. de Laat, P. G. de Groot, Autoantibodies directed against domain I of beta2-glycoprotein I. Curr Rheumatol Rep 13, 70–76 (2011).

21. G. M. Iverson, E. J. Victoria, D. M. Marquis, Anti-beta2 glycoprotein I (beta2GPI) autoantibodies recognize an epitope on the first domain of beta2GPI. Proc Natl Acad Sci U S A 95, 15542–15546 (1998).

22. A. Banzato et al., Antibodies to Domain I of beta(2)Glycoprotein I are in close relation to patients risk categories in Antiphospholipid Syndrome (APS). Thromb Res 128, 583–586 (2011).

23. B. de Laat et al., Immune responses against domain I of beta(2)-glycoprotein I are driven by conformational changes: domain I of beta(2)-glycoprotein I harbors a cryptic immunogenic epitope. Arthritis Rheum 63, 3960–3968 (2011).

24. B. Bouma et al., Adhesion mechanism of human beta(2)-glycoprotein I to phospholipids based on its crystal structure. EMBO J 18, 5166–5174 (1999).

25. R. Schwarzenbacher et al., Crystal structure of human beta2-glycoprotein I: implications for phospholipid binding and the antiphospholipid syndrome. EMBO J 18, 6228–6239 (1999).

26. M. Hammel et al., Solution structure of human and bovine beta(2)-glycoprotein I revealed by small-angle X-ray scattering. J Mol Biol 321, 85–97 (2002).

27. C. Agar et al., Beta2-glycoprotein I can exist in 2 conformations: implications for our understanding of the antiphospholipid syndrome. Blood 116, 1336–1343 (2010).

28. I. Buchholz, P. Nestler, S. Koppen, M. Delcea, Lysine residues control the conformational dynamics of beta 2-glycoprotein I. Phys Chem Chem Phys 20, 26819–26829 (2018).

29. C. Agostinis et al., In vivo distribution of beta2 glycoprotein I under various pathophysiologic conditions. Blood 118, 4231–4238 (2011).

30. S. S. Pierangeli et al., Toll-like receptor and antiphospholipid mediated thrombosis: in vivo studies. Ann Rheum Dis 66, 1327–1333 (2007).

31. R. T. Urbanus, M. T. Pennings, R. H. Derksen, P. G. de Groot, Platelet activation by dimeric beta2-glycoprotein I requires signaling via both glycoprotein Ibalpha and apolipoprotein E receptor 2’. J Thromb Haemost 6, 1405–1412 (2008).

32. S. Ramesh et al., Antiphospholipid antibodies promote leukocyte-endothelial cell adhesion and thrombosis in mice by antagonizing eNOS via beta2GPI and apoER2. J Clin Invest 121, 120–131 (2011).

33. J. Zhang, K. R. McCrae, Annexin A2 mediates endothelial cell activation by antiphospholipid/anti-beta2 glycoprotein I antibodies. Blood 105, 1964–1969 (2005).

34. Z. Romay-Penabad et al., Apolipoprotein E receptor 2 is involved in the thrombotic complications in a murine model of the antiphospholipid syndrome. Blood 117, 1408–1414 (2011).

35. M. Chinnaraj, W. Planer, V. Pengo, N. Pozzi, Discovery and characterization of 2 novel subpopulations of aPS/PT antibodies in patients at high risk of thrombosis. Blood Adv 3, 1738–1749 (2019).

36. M. Chinnaraj et al., Structure of prothrombin in the closed form reveals new details on the mechanism of activation. Sci Rep 8, 2945 (2018).

37. M. Chinnaraj, W. Planer, N. Pozzi, Structure of Coagulation Factor II: Molecular Mechanism of Thrombin Generation and Development of Next-Generation Anticoagulants. Front Med (Lausanne) 5, 281 (2018).

38. A. I. Okemefuna, R. Nan, J. Gor, S. J. Perkins, Electrostatic interactions contribute to the folded-back conformation of wild type human factor H. J Mol Biol 391, 98–118 (2009).

39. E. M. Bevers, R. F. Zwaal, G. M. Willems, The effect of phospholipids on the formation of immune complexes between autoantibodies and beta2-glycoprotein I or prothrombin. Clin Immunol 112, 150–160 (2004).

40. V. Pengo et al., Lupus anticoagulant identifies two distinct groups of patients with different antibody patterns. Thromb Res 172, 172–178 (2018).

41. N. Pozzi, D. Bystranowska, X. Zuo, E. Di Cera, Structural Architecture of Prothrombin in Solution Revealed by Single Molecule Spectroscopy. J Biol Chem 10.1074/jbc.M116.738310 (2016).

42. E. Lerner et al., Toward dynamic structural biology: Two decades of single-molecule Forster resonance energy transfer. Science 359 (2018).

43. F. H. Passam et al., Beta 2 glycoprotein I is a substrate of thiol oxidoreductases. Blood 116, 1995–1997 (2010).

44. I. V. Gopich, A. Szabo, Single-molecule FRET with diffusion and conformational dynamics. J Phys Chem B 111, 12925–12932 (2007).

45. A. G. Kikhney, D. I. Svergun, A practical guide to small angle X-ray scattering (SAXS) of flexible and intrinsically disordered proteins. FEBS Lett 589, 2570–2577 (2015).

46. I. Dienava-Verdoold et al., Patient-derived monoclonal antibodies directed towards beta2 glycoprotein-1 display lupus anticoagulant activity. J Thromb Haemost 9, 738–747 (2011).

47. J. E. Hunt, R. J. Simpson, S. A. Krilis, Identification of a region of beta 2-glycoprotein I critical for lipid binding and anti-cardiolipin antibody cofactor activity. Proc Natl Acad Sci U S A 90, 2141–2145 (1993).

48. Y. Sheng, D. A. Kandiah, S. A. Krilis, Anti-beta 2-glycoprotein I autoantibodies from patients with the “antiphospholipid” syndrome bind to beta 2-glycoprotein I with low affinity: dimerization of beta 2-glycoprotein I induces a significant increase in anti-beta 2-glycoprotein I antibody affinity. J Immunol 161, 2038–2043 (1998).

49. M. Hammel et al., Mechanism of the interaction of beta(2)-glycoprotein I with negatively charged phospholipid membranes. Biochemistry 40, 14173–14181 (2001).

50. Y. Ioannou et al., Binding of antiphospholipid antibodies to discontinuous epitopes on domain I of human beta(2)-glycoprotein I: mutation studies including residues R39 to R43. Arthritis Rheum 56, 280–290 (2007).

51. G. M. Willems et al., Role of divalency in the high-affinity binding of anticardiolipin antibody-beta 2-glycoprotein I complexes to lipid membranes. Biochemistry 35, 13833–13842 (1996).

52. A. Banzato et al., Circulating beta2 glycoprotein I-IgG anti-beta2 glycoprotein I immunocomplexes in patients with definite antiphospholipid syndrome. Lupus 21, 784–786 (2012).

53. P. L. Meroni, M. O. Borghi, E. Raschi, F. Tedesco, Pathogenesis of antiphospholipid syndrome: understanding the antibodies. Nat Rev Rheumatol 7, 330–339 (2011).

54. C. J. Lee, A. De Biasio, N. Beglova, Mode of interaction between beta2GPI and lipoprotein receptors suggests mutually exclusive binding of beta2GPI to the receptors and anionic phospholipids. Structure 18, 366–376 (2010).

55. A. Kolyada, D. A. Barrios, N. Beglova, Dimerized Domain V of Beta2-Glycoprotein I Is Sufficient to Upregulate Procoagulant Activity in PMA-Treated U937 Monocytes and Require Intact Residues in Two Phospholipid-Binding Loops. Antibodies (Basel) 6 (2017).

56. T. McDonnell et al., Development of a high yield expression and purification system for Domain I of Beta-2-glycoprotein I for the treatment of APS. BMC Biotechnol 15, 104 (2015).

57. N. Pozzi et al., Chemical synthesis and characterization of wild-type and biotinylated N-terminal domain 1-64 of beta2-glycoprotein I. Protein Sci 19, 1065–1078 (2010).

58. T. C. R. McDonnell et al., PEGylated Domain I of Beta-2-Glycoprotein I Inhibits the Binding, Coagulopathic, and Thrombogenic Properties of IgG From Patients With the Antiphospholipid Syndrome. Front Immunol 9, 2413 (2018).

59. P. Durigutto et al., New insight into antiphospholipid syndrome: antibodies to beta2glycoprotein I-domain 5 fail to induce thrombi in rats. Haematologica 10.3324/haematol.2018.198119 (2018).

60. A. Kolyada, A. Porter, N. Beglova, Inhibition of thrombotic properties of persistent autoimmune anti-beta2GPI antibodies in the mouse model of antiphospholipid syndrome. Blood 123, 1090–1097 (2014).

61. Z. Otwinowski, W. Minor, Processing of x-ray diffraction data collected by oscillation methods. Methods Enzymol. 276, 307–326 (1997).

62. E. J. Dodson, M. Winn, A. Ralph, Collaborative Computational Project, number 4: providing programs for protein crystallography. Methods Enzymol 277, 620–633 (1997).

63. P. Emsley, K. Cowtan, Coot: model-building tools for molecular graphics. Acta Crystallogr D Biol Crystallogr 60, 2126–2132 (2004).

64. J. Lozier, N. Takahashi, F. W. Putnam, Complete amino acid sequence of human plasma beta 2-glycoprotein I. Proc Natl Acad Sci U S A 81, 3640–3644 (1984).

65. J. Hunt, S. Krilis, The fifth domain of beta 2-glycoprotein I contains a phospholipid binding site (Cys281-Cys288) and a region recognized by anticardiolipin antibodies. J Immunol 152, 653–659 (1994).

66. L. Acquasaliente et al., Molecular mapping of alpha-thrombin (alphaT)/beta2-glycoprotein I (beta2GpI) interaction reveals how beta2GpI affects alphaT functions. Biochem J 473, 4629–4650 (2016).

67. A. Kolyada, A. De Biasio, N. Beglova, Identification of the binding site for fondaparinux on Beta2-glycoprotein I. Biochim Biophys Acta 1834, 2080–2088 (2013).

